# Incidence, antifungal resistance properties, and virulence traits of candida species isolated from HIV/AIDS Patients from the hospital system in Kenya

**DOI:** 10.1101/2022.12.08.519702

**Authors:** Haron N. Miruka, Omwenga O. Eric, Stanslaus Musyoki, Silas O. Awuor

**Affiliations:** School of Health Sciences, Kisii University P.O BOX 408-40200 Kisii, Kenya; Department of Medical Microbiology & Parasitology, School of Health Sciences, Kisii University P.O. Box 408 – 40200 Kisii; Department of Medical Laboratory Sciences, School of Health Sciences, South Eastern Kenya University; P.O. Box 170-90200, Kitui, Kenya; Microbiology department, Jaramogi Oginga Odinga Teaching and Referral Hospital, P.O Box 849, Kisumu, Kenya

**Keywords:** Antifungal Susceptibility, Resistance genes, Antibiofilm activity, Virulence traits, Candida Specie, HIV

## Abstract

Candidiasis is the most common fungal infection in hospitalized patients with acquired immune deficiency syndrome resulting to morbidity and mortality. This study aimed at characterizing, incidence, susceptibility, resistance genes, antibiofilm activity, and virulence traits of *Candida* species isolated from HIV-Infected patients. One hundred and eighty-one samples were collected and cultured on Sabouraud Dextrose Agar, biochemical tests and confirmed using automated Vitek-2 ^®^ Compact bioMérieux followed by susceptibility tests. were done by use of various conventional antifungals against the isolates using standard procedures. Virulence factors, biofilm formations and resistance genes of Candida strains were determined. Out of the 181 samples, 46 were identified as *Candida spp*., 20 *C. albicans* (43.5%), 6 *C. tropicalis* (13.0%), 8 *C. krusei* (17.4%), 4 *C. glabrata* (8.7%), 3 *C. famata* (6.5%), 3 *C. parapsilosis* (6.5%), and *2 C. guilliermondii* (4.3%). All the *Candida albicans* isolated were both Gram positive and Germ test tube test positive. Eighteen (90%) of the isolates were susceptible to Clotrimazole at a concentration of 5 μg/mL – 10μg/mL followed by 17 (85%) isolates to Panosoconazole at a concentration of 0.002 μg/mL – 5μg/mL. Eight (40.0 %) of the *Candida albicans* isolates possessed the gene (*cdr1*) that was observed at 286 bp. Virulence enzymes was determined in which 100% produced Haemolysin, followed by proteinase (75.0%), phospholipase (50%), coagulase at (50%) and lastly capsulase (25.0%). Fluconazole and Clotrimazole did not inhibit growth of *C. albicans* at high concentrations but from our study, it was deduced that they inhibit biofilm formation at lower concentrations. *C. albicans* isolates were resistant to multiple antifungal including those commonly used in the management on HIV/AIDs patient. This attributed to resistant genes and produced various virulence factors that were found to be present in the isolates. Therefore, there is a need to carry out regular surveillance on antifungal drug resistance.

## Introduction

The decline of CD4^+^ T lymphocyte count in HIV patients invites the risk of acquired immunodeficiency syndrome (AIDS) (1). Though the introduction of antiretroviral therapy has a major impact, candidiasis still remains a common opportunistic infection. Increased infections, prolonged use of antifungals to treat recurrent infections, and the emergence of antifungal resistance have created the need for antifungal susceptibility testing (2). The knowledge about the effects and burden of antifungal resistance is less compared to that of antibiotic-resistant bacterial infections, which are widely recognized as a public health problem (3). This highlights the need to understand the reasons for their emergence, create awareness among medical and public health communities about these infections, and greater attention to prevent and control them.

*Candida* species are the most common cause of fungal infections worldwide. *Candida* species are normal microbiota within the gastrointestinal tracts, respiratory tracts, vaginal area and mouth. Additionally, they have been classified as one of the most commonly sort sexually transmitted diseases amongst sexually active individuals (4). *Candida* is a yeast growth present in all females and is normally controlled by bacteria. *Candida* species differ in their antifungal susceptibility and virulence factors. The genus is composed of a heterogeneous group of organisms, and more than 17 different *Candida* species are known to be etiological agents of human infections; however, more than 90% of invasive infections are caused by *C. albicans, C. glabrata, C. parapsilosis, C. tropicalis, C. krusei, C. dubliniensis*, and *C. lusitaniae* (5). The yeast begins to invade and colonize the body tissues by releasing toxins like killer toxins which are protein in nature which kill sensitive cells of the same or related yeast genera into the bloodstream causing symptoms like: lethargy, chronic diarrhea, yeast vaginitis, bladder infections, muscle and joint pain, menstrual problems, constipation and severe depression [6, 7, 8].The opportunistic fungi that most commonly affect humankind are the species of *Candida* and *Cryptococcus* (8), out of these two groups, the *Candida albicans* species remains the most predominant *Candida* species causing candida infections with cases that numbers to more than a half in the world (6). An increase in the prevalence of yeast infections caused by non-albicans *Candida* such as *Candida tropicalis, Candida glabrata, Candida parapsilosis*, and *Candida krusei*, have also been reported in some parts of the world (8)

Prolonged usage of antifungals in treating infections caused by *C. albicans* has led to the emergence of azole resistance. This acquired azole resistance in clinical isolates of *C. albicans* mostly results in cross-resistance to many unrelated drugs, a phenomenon termed multidrug resistance (MDR) [8, 10, 11].

Fungal etiology of some infections is a reality more and more common in medical practice, facilitated, because of the excessive use of antibiotics, often given in inadequate condition of sickness and taken in excessive doses, and also to the “abuse” of steroids, plus the high incidence of serious debilitating diseases (cancers, HIV infection, diabetes) which requires aggressive therapy or combination therapies that may have a negative effect by decreasing the overall body strength [12,13]. Some *Candida* spp strains on the other hand have been documented to form biofilms which is currently becoming a big problem in management of fungal pathogens worldwide. Most of the candida spp are capable of producing other virulence factors (phospholipase, protease, hemolysin, coagulase factors etc.) which help the fungi to survive *in vivo*, penetration to the mucous layer and adherence to the underlying epithelial cells, which enables them to evade the host defense system. Therefore, this study aimed at characterizing, the incidence, susceptibility patterns, resistance genes, antibiofilm activity, and virulence traits of *Candida* species isolated from HIV/AIDS Patients Attending at a Kenyan medical facility-Muhoroni county Hospital.

## Materials and methods

### i. Study design

This was a cross-sectional study conducted between January to August 2020 in the department of Comprehensive Care Centre (CCC), Muhoroni sub-county Hospital (MCH) which is a consultant and a referral center for the sub county. *Candida spp*. isolated from patients suffering from oral thrush, vaginal candidiasis, candiduria, oesophageal candidiasis, and candidemia were included and characterized. Candidemia was considered as invasive candidiasis, while oral thrush, vaginal candidiasis, oesophageal candidiasis, and candiduria were regarded as non-invasive candidiasis.

### ii. Study area

The study was conducted at Muhoroni County Hospital (0.15660° S, 35.19840º E) in Kisumu County (East to Muhoroni Sugar Company), Kenya. Muhoroni County Hospital is a Government 100-bed capacity health facility located in Muhoroni Town, Muhoroni Sub County in Kisumu County.

### iii. Study Population

### iv. Study samples

This study targeted total HIV infected patients with low detectable viral load (LDL)who sought Medicare at Muhoroni County Hospital (MCH) between January to August 2020 showing the symptoms of Candidiasis. Based on Harris et al. (1991) formulae a sample size of 181 sample size was utilized in this study. We targeted various clinical samples which included high vaginal swabs (HVS) from female patient, urine and sputum from both female and male patients seeking medicare at the comprehensive care center department within the study period. The samples were then kept in the laboratory under −18°C before the analysis.

### v. Laboratory analysis

#### a. Candida spp. Identification and Preparation of Inoculums

The primary isolation of yeasts was performed using the CHROMagar ® Candida (Difco) Chromogenic differential medium in Petri dishes and incubated at 37°C for 48h using protocols that have been used previously [14]. Green colored colonies were identified as *C. albicans*, blue-cobalt as *C. tropicalis*, pink or lilac as *C. krusei*, and other species were whitish-pink in color. The isolated yeasts from chromogenic medium were picked and incubated at 35°C for 48 h in tubes with SDA (Sabouraud Dextrose Agar with chloramphenicol – Acumedia) medium and then stored at −20°C for use in the study. The biochemical identification of the yeasts was performed using an automated method (Vitek-2 Compact bioMérieux, Marcy-l’Étoile, France). The Biomerieux Vitek-2 system includes the Vitek2 cards that allow species identification by comparison of the biochemical profile with an extensive database Biomerieux Vitek-2 expanded its role in this area with a yeast susceptibility test that determines Candida growth spectrophotometrically using Vitek-2 microbiology systems, performing fully automated testing of susceptibility to 5-FC, AMB, FCZ, and VCZ (15). For the preparation of the fungal inoculum (3 mL 0.45% saline + yeast colony), a McFarland scale 2 from the DensiChek-bioMerieuxVitek system was used. This standardized suspension was aspirated into the identification cards, and then the cards were sealed and subjected to biochemical tests by an optical sensor reading. We used the YST card (Yeast identification, bioMérieux) to determine the genus and species of yeast. The test was considered complete when the percentage of probability was ≥85% and there was no request for further testing.

##### Gram staining

Smears were prepared from growths that appeared on the SDA (Oxoid, UK) allowed to air-dry, heat-fixed and then stained by the Gram staining method and examined microscopically as per a previously used protocol (16).

##### Germ tube test

All yeasts isolated from the specimens processed were identified to the species level using the germ tube test and API ID 32 C strip (bioMerieux, Marcy-l’Etoile, France) (14). Briefly, a pasteur pipette was used to collect 5.0ml of fresh human serum (from the serology laboratory at the Muhoroni County Hospital) then put into labeled test tubes. Using a sterile inoculating loop, a colony of yeasts was transferred into the serum in the labeled test tubes. The colony was then gently emulsified in the serum. Incubated at 37°C for about 3 hours. Using a Pasteur pipette, a drop of the suspension was taken from the test tube after incubation and placed on a clean dry slide for examination. The suspension was covered with a coverslip and examined under a microscopically under low power (10X objective lens) for germ tubes on the yeasts. High power objective (40X objective) was used to confirm the presence or absence of germ tubes. The Presence of short hyphal (filamentous) extension arising from a yeast cell indicated the presence of *C. albicans* (16). Both the positive control isolates (*C. albicans* - ATCC 10231) and the negative control isolate (*C. glabrata* -ATCC 2001) were included in this test. All tests were done in triplicates independent of each other.

##### API ID 32 C kits

API ID 32 C kits were used to deduce for the presence of *C. albicans* and to identify the yeast isolates as used before (17). Briefly, by use of a sterile inoculating loop, one or several colonies from the Sabouraud’s dextrose agar (Oxoid, UK) plates was transferred into 2 ml of API® suspension medium (bioMerieux, Marcy-l’Etoile, France). The colonies were emulsified in the API® suspension medium to form a suspension of turbidity equivalent to 2 McFarland.

The turbidity of the suspension was measured with the Densimat (bioMerieux, Marcy-l’Etoile, France) device which accompanied the test kit. 250μl of the suspension (already prepared) were then transferred into 7ml of API C medium (bioMerieux, Marcy-l’Etoile, France) using ATB electronic pipette (bioMerieux, Marcy-l’Etoile, France) supplied with the test kits. All tests were done in triplicates that are independent of each other.

#### b. In vitro Antifungal Susceptibility Tests

Thereafter, antifungal susceptibility tests (AFST) were conducted using the ViteK-2^®^ automated system and Etest^®^ (Biodisk AB, Solna, Sweden) according to the manufacturer’s recommendations. These methods have been chosen because they are easy to perform and offer results in a short period of time (18).

Briefly, 180 μL of inoculum was standardized to the McFarland scale 2.0 using the DensiChek densitometer of the ViteK-2^®^ system, placed in a tube containing 3 mL of 0.45% saline, and aspirated into the AST-YSO1 card (bioMérieux). The following antifungal drugs were tested: Amphotericin B (10 μg/), Fluconazole (25 μg), Itraconazole (10 μg), Nystatin (50 μg), Clotrimazole (10 μg), and Panosoconazole (5 μg). The analysis and interpretation of data were performed according to M27-A3 and M27-S3 CLSI standards. For quality control, *C. krusei* (ATCC 6258) and *C. parapsilosis* (ATCC 22019) were used as standard strains. For fungal strains that did not respond to the AST cards, an Etest^®^ (Biodisk AB, Solna, Sweden) was performed, which consisted of a gradient method with predefined concentrations of FCZ and AMB in μg/mL. In this method, the colonies were seeded at a concentration of 0.5 McFarland on dishes with RPMI (Probac^®^, São Paulo, Brazil) supplemented with 2% glucose agar and then incubated at 35°C for 48 h. The reading was performed by assessing the point of intersection between the halo formed and the Etest strip. The Fluconazole MIC was determined as the lowest concentration that inhibited 80% of fungal strains, and the Amphotericin MIC was calculated as the lowest concentration with no observed fungal growth. The profile of the antifungal drug sensitivity was classified as sensitive (S), dose-dependent sensitivity (S-DD), and/or intermediate (I), and resistant (R).

The breakpoints used to define sensitivity, intermediate, and resistance for each species were those defined by Clinical and Laboratory Standards Institute [CLSI] (2022). The MIC values ≤ 8 μg/mL for Fluconazole were considered susceptible (S), 16– 32 μg/mL was considered as susceptible dose-dependent (SDD), and ≥64 μg/mL as resistant (R). For Amphotericin, MICs ≤ 1 μg/mL were considered to be S and ≥1 μg/mL was R. For Itraconazole, CIMs ≤ 4 μg/mL were considered to be S, 8–16 μg/mL was I, and ≥32 μg/mL was R. For Panosoconazole, MICs ≤ 0,125 μg/mL were S and ≥16 μg/mL were considered R. Fluconazole and amphotericin were chosen for the tests because they have different mechanisms of action and are the main drugs chosen for the treatment of Candida infections (19). Panosoconazole and Itraconazole were chosen because they could be an alternative for species resistant to Fluconazole and Amphotericin.

#### c. Determination of biofilm formation by microtitration plate method

After the Candida spp. isolates produced in the SDB were adjusted to be 10^7^ cfu / mL, they were distributed as 20 μL in each well of the 96-well plate for purposes of determining biofilm formation in presence of antifungals using previously used and established protocol (23. Briefly, 180 μL synthetic dextrose liquid (SDL) medium containing 2.5% glucose was transferred onto it and incubated for 48 h at 35 ° C. After incubation, the plates were emptied and each well was washed 3 times with sterile physiological saline. The wells were fixed with 200μL 99% methanol for 15 minutes. At the end of this period, the wells were emptied and left to dry. Subsequently, each well was stained with 200 μL 2% crystal violet for 15 minutes. When this period ended, the wells were washed with distilled water and dried for 1hr. After the drying, the wells were treated with 160μL 33% glacial acetic acid and assessed spectrophotometrically at 590 nm. For each experiment, background staining was corrected by subtracting the crystal violet bound to un-treated controls (Blank) from those of the tested sample. The experiments were done in triplicate and average OD_590_nm values were calculated. To estimate the antibiofilm activity (Abf A) of a given antifungal the following equation will be used Abf A (%) = (1-(OD_Test sample_ - OD_Blank_)/ (OD _Untreated sample_ – OD _Blank_) × 100.

#### d. Genome Characterization for resistance genes

##### Polymerase Chain Reaction (PCR) for pathogenic and antibiotic resistant genes

DNA was extracted from the isolates using a method described by (20). The DNA for the resistant isolates were amplified using conventional PCR as used before (22) and were used in detecting the presence of *CDR1, CDR2, RTL3* and *MAL 2* resistance genes amongst the isolates. The list of the primers used are presented in **table 2.1** below. The cycles were set at 94-98ºC for 1-3 minutes for denaturing, depending on the DNA template, 55-72ºC. for 1-2 minutes for annealing and extension at a temperature of 68-72ºC depending on DNA polymerase rate and length of the target DNA.

**Table 2.1:**
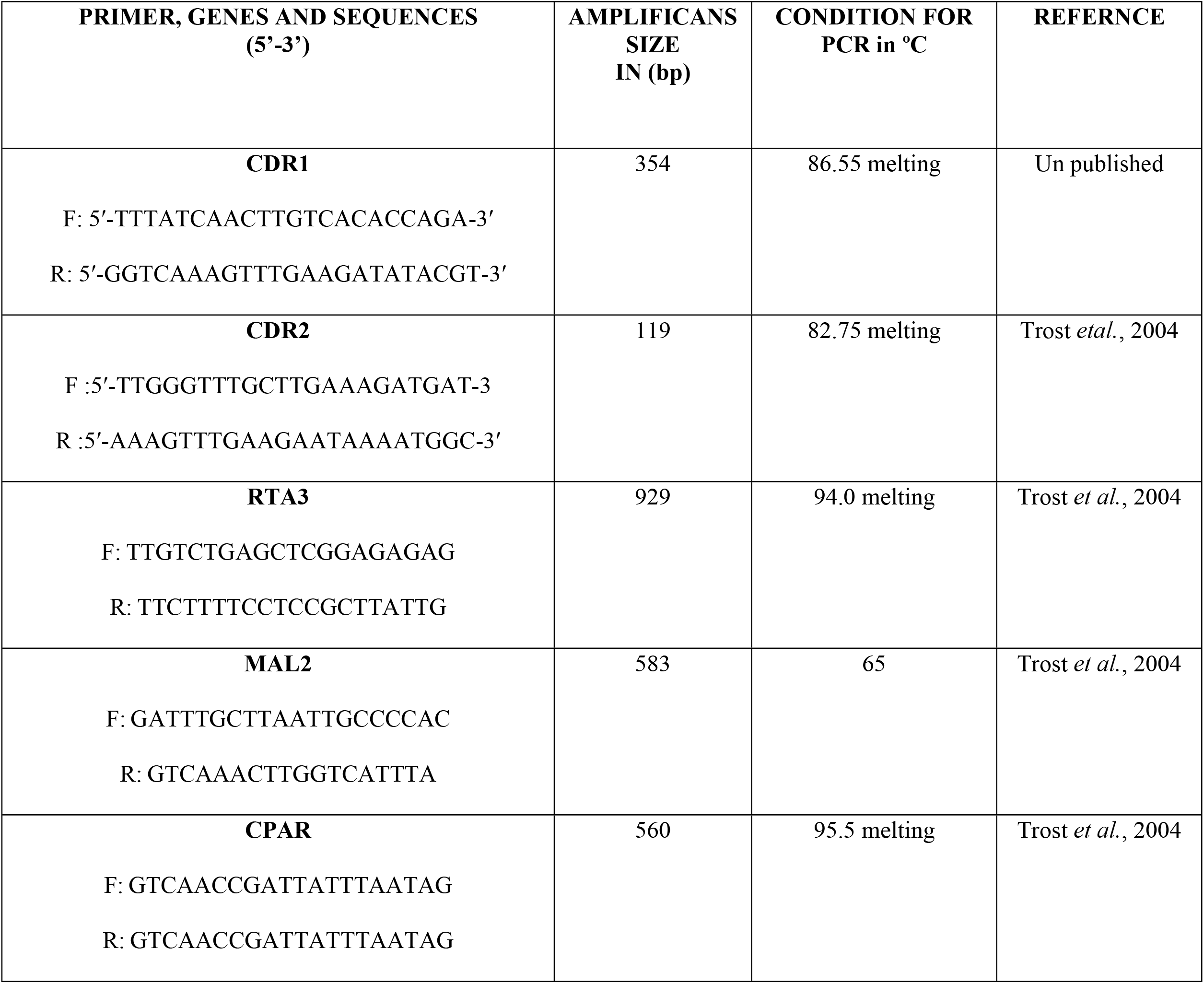
Sequences, conditions and amplificons of oligonucleotide primers in PCR used for the study

##### Gel electrophoresis

The amplified candida spp. isolates genome was separated by use of PFGE as used before (18) with some slight changes. A two-block program with a first block ramp time of 2 sec to 10 sec for 13 hrs. at 6V/cm (for separation of smaller fragments) and second block with ramp time of 20 and 25 sec for 6 hrs. (for separation of larger fragments) at 6V/cm were used. The NotI restriction enzyme was used to digest the chromosomal DNA. Restriction fragments were separated in a 1% pulsed-field certified agarose gel in 1× TBE (8.9 mM Tris base, 8.9 mM boric acid, and 0.25 mM disodium EDTA) by using a CHEF-DR II system (Bio-Rad). CHEF DNA size standard of *Saccharomyces cerevisiae* (Bio-Rad Laboratories, Inc, CA) was then used as a molecular mass standard. Following electrophoresis, gels were stained for 20 min with ethidium bromide (2μg/ml in 1% TBE buffer), distained.

#### e. Detection of other virulence factors

##### Extracellular Phospholipase Activity

Phospholipases were screened for by measuring the size of the zone of precipitation after growth of the isolates on egg yolk agar using established methods as used before (13). The plates were read with the aid of a computerized image analysis system (Quantimet 500 Qwin; Leica), which measures the diameter of the colonies relative to the precipitation zones on a magnified scale. Phospholipase activity was expressed as the ratio of the diameter of the colony to the diameter of the colony plus the precipitation zone (in mm) (20). All tests will be done in triplicates that are independent of each other.

##### Determination of Haemolysin production activity

The β-haemolytic activity was tested for on base agar (Himedia, India) supplemented with 7 % sheep erythrocytes for 18–24 h as described by (38). Briefly, one colony of the overnight culture was emulsified in 1ml of sterile normal saline then 10 μl of suspension was inoculated onto an SDA plate supplemented with 3% glucose and 7% sheep blood. Plates were then incubated aerobically at 37°C for 72 hours. Haemolysin activities were determined by measuring the diameter of the translucent halo around the inoculum site when viewed with transmitted light. *Streptococcus pyogenes* ATCC12384 was used as the positive control. All tests were done in triplicates that are independent of each other.

##### Determination of coagulase activity

Coagulase activities were detected and interpreted using coagulase biochemical test as described previously (22). Briefly, Candida cells were inoculated into Sabouraud’s dextrose broth and incubated aerobically at 37°C for 18–24 hours. A total of 100 μl of the overnight broth were aseptically placed into a test tube containing 0.5 ml of EDTA rabbit plasma and incubated aerobically at 37°C for 4 hours. *Staphylococcus aureus* ATCC 25923 was used as positive control while *Staphylococcus epidermidis* ATCC 14990 was used as the negative control. All tests were done in triplicates that are independent of each other.

##### Determination of capsulase formation ability

The evaluation of capsule production was assessed by culturing the candida strains on Congo red agar (CRA) as described by (23) with minor modifications. The Congo red agar was prepared by mixing 36 g saccharose (Sigma Chemical Company, Lezennes, France) with 0.8 g Congo red in 1 L of on Tryptic Soy Agar (TSA, Difco, City, Spain) supplemented with NaCl (1.0%). The plates were then inoculated with the Candida spp. and incubated aerobically for 24 h at 37ºC followed by another 24 h at 30°C. After incubation, black colonies were considered as capsule producers, whereas red colonies were considered as non-producers (23). All tests were done in triplicates that are independent of each other.

##### Determination of Proteinase Production Activity

Extracellular protease activity was assessed by plating the Candida isolates on bovine serum albumin agar (BSA) with 1% KH_2_PO_4_, 0.05% MgSO_4_ (LOBACHEMIE-Mumbai, India), 2% agar, 0.01% yeast extract (Finkem), and 0.2% BSA (AppliChem, Germany) adjusted to pH of 4.5 as previously described (22). Clearance zone around the colonies as result of the cleavage of the BSA by proteases was indicative of the positive protease activities. The diameter clearance was measured and recorded after 72 h of growth. All tests were done in triplicates that are independent of each other.

##### Validity and reliability

All experiments were conducted in triplicates that were independent of each other to validate reproducibility.

## Data analysis

Statistical analysis was performed using Stata Software. Data on socio-demographics were summarized by frequencies and percentages. All values of diameter zones of inhibition are reported as mean ± standard error.

## Ethical consideration

Confidentiality and privacy were strictly adhered to and no names of individuals were recorded or made known in the collection or reporting of information. The study was granted ethical clearance by the Board of Postgraduate Studies (BPS) of Kisii University Ref no. KSU/R&E/03/5/513(**Supplementary data 1**), and ethical approval to conduct the study was sought from the Institutional Research Ethics Committee (IREC) at Moi University/Moi Teaching and Referral Hospital (MTRH) Ref. No. IREC/2019/111(**Supplementary data 2**) and the National Commission of Science, Technology and Innovations (NACOSTI) Ref. No. NACOSTI/P/20/3219 (**Supplementary data 3**).

## Results

### i. Demographics and Incidence of Isolates

From 181 AIDS patients participating in the study, 46 (25.4%) fungal isolates were recovered from these patients, and thus, in some cases, more than one species was isolated from a single clinical sample. Eighty-three percent (83%) of patients participating in the study had oropharyngeal candidiasis, and 52.2% had used antifungals. Of these, 75% were using both Fluconazole and nystatin and 8.3% used Fluconazole, nystatin, and Amphotericin. Among the samples, 46 (25.4%) grew in CHROMagar Candida medium and 135(74.6%) were negative. Using the automated Vitek-2 ^®^ Compact bioMérieux system, 46 isolates were identified as Candida spp., that included 20 *C. albicans* (43.5%), 6 *C. tropicalis* (13.0%), 8 *C. krusei* (17.4%), 4 *C. glabrata* (8.7%), 3 *C. famata* (6.5%), 3 *C. parapsilosis* (6.5%), and 2 *C. guilliermondii* (4.3%) (Table 3.1). When looking at the distribution of isolates of Candida spp. by patients’ gender, we observed that the number of isolates coming from the women patients 40 (87%) were higher than that from the men 6 (13%) as shown in Table 3.1.

**Table 3.1:** Number and incidence of the Candida strains identified by the Vitek-2^®^ system, and their distribution according to patient gender.

| Species | Female | Male | Total No. of isolates |
| --- | --- | --- | --- |
| *Candida albicans* | 20 (43.5%) | 0 (0%) | 20 |
| *Candida tropicalis* | 5 (4.3%) | 1 (2.2%) | 6 |
| *Candida krusei* | 6 (13.0%) | 2 (4.3%) | 8 |
| *Candida glabrata* | 2 (4.3%) | 2 (4.3%) | 4 |
| *Candida guilliermondii* | 2 (4.3%) | 0 (0%) | 2 |
| *Candida parapsilosis* | 2 (4.3%) | 1 (2.2%) | 3 |
| *Candida famata* | 3 (6.5%) | 0 (0%) | 3 |
| **Total** | **40 (87%)** | **6 (13%)** | **46** |

### ii. Characterization of *C. albicans*

Of the 20 *Candida albicans* isolated, Gram staining technique, Germ test tube and sub culture on SDA media (**Supplementary data 6**) was performed for confirmation of the species of interest and they both revealed that 100% of the isolates were *Candida albicans* as shown in Table 3.2. For instance, in Gram staining all the isolates were found to be Gram positive in cocci as shown in supplementary data S 4. Also, for germ tube test the 20 isolates proved to be positive by producing yeast cell budding like shapes as shown in supplementary data 5. Regarding culturing on SDA media, the isolates produced white to cream with smooth & yeast like appearance colonies (Supplementary data 6).

**Table 3.2:** Morphological characteristics of isolated organism of *C. albicans*

| VARIABLE | Frequency<br>(N=20) | Percentage<br>(%) |
| --- | --- | --- |
| **Gram Staining** |  |  |
| • Positive | 20 | 100% |
| • Negative | 00 | 00% |
| **Germ Test Tube** |  |  |
| • Positive | 20 | 100% |
| • Negative | 00 | 00% |
| **Colony Appearance on SDA Media** |  |  |
| • White to cream with smooth & yeast like appearance | 20 | 100% |
| • Non- White to cream with smooth & yeast like appearance | 00 | 00% |

The isolates were further characterized using **API ID 32 C** kits and all the 20 isolates were confirmed to be *C. albicans* by turning out to be positive to SAC, RAF, TRH, ROR and RHA and Negative to ARA, CEL MAL MEL and MAN as shown in table 3.3

**Table 3.** Biochemical characterization of the isolate to API ID 32 C kits.

| ISOLATE | API 32C KIT |  |  |  |  |  |  |  |  |  |
| --- | --- | --- | --- | --- | --- | --- | --- | --- | --- | --- |
|  | SAC | ARA | CEL | RAF | MAL | TRH | SOR | RHA | MEL | MAN |
| MCH/05/20 | + | - | - | + | - | + | + | + | - | - |
| MCH/06/20 | + | - | - | + | - | + | + | + | - | - |
| MCH/07/20 | + | - | - | + | - | + | + | + | - | - |
| MCH/13/20 | + | - | - | + | - | + | + | + | - | - |
| MCH/14/20 | + | - | - | + | - | + | + | + | - | - |
| MCH/16/20 | + | - | - | + | - | + | + | + | - | - |
| MCH/33/20 | + | - | - | + | - | + | + | + | - | - |
| MCH/34/20 | + | - | - | + | - | + | + | + | - | - |
| MCH/36/20 | + | - | - | + | - | + | + | + | - | - |
| MCH/37/20 | + | - | - | + | - | + | + | + | - | - |
| MCH/38/20 | + | - | - | + | - | + | + | + | - | - |
| MCH/39/20 | + | - | - | + | - | + | + | + | - | - |
| MCH/47/20 | + | - | - | + | - | + | + | + | - | - |
| MCH/62/20 | + | - | - | + | - | + | + | + | - | - |
| MCH/67/20 | + | - | - | + | - | + | + | + | - | - |
| MCH/74/20 | + | + | + | - | + | + | + | + | + | + |
| MCH/75/20 | + | + | + | + | + | + | + | + | + | + |
| MCH/82/20 | + | + | - | + | + | + | + | - | + | + |
| MCH/123/20 | + | + | + | + | + | + | + | + | + | + |

**KEY:** SAC = D –Saccharose, ARA = L – Arabinose, CEL = D –Cellobiose, RAF = D – Rafimose, MAL = D – Maltose, TRF = D- Trihalose, SOR = D – Sorbitol, RHA = L – Rhamnose, MEL = D –Mellobiose, MAN = D –Manitol, +=Positive and -= Negative.

### iii. Distribution of *C. albicans* among the age and gender in the study sites

The age group with the highest frequency of *Candida albicans* isolates was 16–25 years at 8 (40%), followed by age group of 26–35 years at 7 (35%), age group 5-15 years at 4 (20%), age group 36-45 years at 1 (5%) and lastly age group >46 years at 0 (0%). From the samples analyzed there were high isolates on HVS samples n=12/76 followed by urine sample (n=8/90) and lastly sputum samples at 0/15 as show in the table 3.4.

**Table 3.4:** distribution of isolated *C. albicans* isolates from the analyzed samples by age groups

| AGE GROUP | URINE SAMPLE(n=90) |  | SPUTUM SAMPLE(n=15) |  | HVS SAMPLE(n=76) |  |
| --- | --- | --- | --- | --- | --- | --- |
|  | Frequency | Percentage | Frequency | Percentage | Frequency | Percentage |
| 5-15 | 3 | 33.3% | 0 | 0% | 1 | 1.3% |
| 16-25 | 2 | 22.2% | 0 | 0% | 6 | 7.9% |
| 26-35 | 3 | 33.3% | 0 | 0% | 4 | 5.3% |
| 36-45 | 0 | 0.0% | 0 | 0% | 1 | 1.3% |
| >46 | 0 | 0.0% | 0 | 0% | 0 | 0% |
| **Total** | **8** |  | **0** |  | **12** |  |

### iv. Antifungal susceptibility patterns

The positive isolates for *C. albicans* were screened for their susceptibility to various antifungal drugs to manage candidiasis cases. Eighteen (90%) isolates were susceptible to Clotrimazole at a concentration of 5 μg/mL – 10μg/mL followed by 17 (85%) isolates to Panosoconazole at a concentration of 0.002 μg/mL – 5μg/mL. However, there was a decreased susceptibility to Fluconazole 16 (80%) which is commonly used for candida management at a concentration of 0.016 μg/mL – 25 μg/mL. In all, 14 (70%) of the *C. albicans* isolates were susceptible to both Amphotericin B and Nystatin at a concentration of 0.016 μg/ mL – 10 μg/mL and 0.016 μg/mL – 50 μg/mL respectively, and *C. albicans* ATCC 10231*-*PC showed 100% susceptibility to almost all antifungal agent used in this study except to Fluconazole and Nystatin which shows 95% both at the above-mentioned concentrations as shown in Table 3.5.

**Table 3.5.** Susceptibility pattern of *C. albicans* isolates against commonly used antifungals drugs to manage candidiasis (n=20)

| ANTIFUNGAL AGENTS | ZONE OF INHIBITION (*C. albicans*) |  |  |  |  |  | ZONE OF INHIBITION (*C. albicans* ATCC 10231-PC) |  |  |  |  |  |
| --- | --- | --- | --- | --- | --- | --- | --- | --- | --- | --- | --- | --- |
|  | S |  | I |  | R |  | S |  | I |  | R |  |
|  | Zone (mm) | N (%) | Zone (mm) | N (%) | Zone (mm) | N (%) | Zone (mm) | N (%) | Zone (mm) | N (%) | Zone (mm) | N (%) |
| FL (25µg) | ≥19 | 16(80) | 18-15 DD | 0 | ≤14 | 4(20) | ≥19 | 19(95) | 18-15 DD | 0 | ≤14 | 1(5) |
| PA (5µg) | ≥17 | 17(85) | 16-14 DD | 0 | ≤13 | 3(15) | ≥17 | 20(100) | 16-14 DD | 0 | ≤13 | 0 |
| IT (10µg) | ≥23 | 10(50) | 22-14 DD | 0 | <13 | 10(50) | ≥23 | 20(100) | 22-14 DD | 0 | <13 | 0 |
| AM (10µg) | ≥15 | 14(70) | 14-10 | 0 | <10 | 6(30) | ≥15 | 20(100) | 14-10 | 0 | <10 | 0 |
| NY (50µg) | ≥15 | 14(70) | 10-14 | 0 | 0 | 6(30) | ≥15 | 19(95) | 10-14 | 0 | 0 | 1(5) |
| CL (10µg) | ≥20 | 18(90) | 12-19 | 0 | ≤11 | 2(10) | ≥20 | 20(100) | 12-19 | 0 | <11 | 0 |

**Key;** FL= Fluconazole, PA= Panosoconazole, IT= Itraconazole, AM= Amphotericin B, NY= Nystatin and CL= Clotrimazole, DD =Disk Diffusion

### v. Detection of other virulence factors

The study investigated the production of various virulence enzymes like proteinase, phospholipase, capsulase formation, coagulase and haemolysin (table 3.6) on the four isolates, which were found to be resistant to all antifungal. It was revealed that 3/4 (75.0 %) of these isolates of *C. albicans* produced proteinase enzyme (supplementary data S7). Also, it was confirmed that two out four isolates (50.0%) produce phospholipases (supplementary data S 8). Further findings also indicate that out of the four isolates two of them (50.0%) had the ability to produce coagulase (supplementary data S 9) and only one isolate (25.0%) produced capsules (supplementary data S 10). Lastly, it was determined that all the four isolates were able to produce the haemolysin by haemolysing the sheep red blood cells causing beta (β) haemolysis (supplementary data S 11). Haemolysin therefore was the most produced virulence factor by these isolates (100%), followed by proteinase (75.0%), phospholipase (50%), coagulase (50%) and lastly capsulase (25.0%). Out of the four isolates one isolates (MCH/16/20) produced all the virulence traits studied, one isolates (MCH/05/20) produced at least three virulence traits and two isolates (MCH/75/20, MCH/47/20) produced at least one of the virulence traits as shown in Table 3.6.

**Table 3.6:** Showing the distribution of the four isolates verses the various virulence traits they produced.

| Isolates | Haemolysis | Proteinase | Phospholipase | Coagulase | Capsulase |
| --- | --- | --- | --- | --- | --- |
| MCH/75/20 | + | - | - | + | - |
| MCH/47/20 | + | + | - | - | - |
| MCH/16/20 | + | + | + | + | + |
| MCH/05/20 | + | + | + | - | - |

**Key;** MCH=Muhoroni County Hospital; (+) = Positive; (-) = Negative

### vi. *C. albicans* isolates resistant genes profiling

Upon screening of the resistant genes, it was deduced that eight (40.0 %) of the *Candida albicans* isolates possessed the gene (*cdr1*) that was observed at 286 bp as shown in Figure 3.1 plate A below. Analysis of the *cdr2* gene revealed that 5 (25.0 %) harbored the *cdr2* gene at 364 bp as shown in Figure 3.1 plate B. It was also revealed that 4 (20.0 %) of the isolates harbored the gene encoded by *rtl3* gene at 201 bp as shown in Figure 3.1 plate C above. The minority, 3 (15.0 %) were confirmed to possess the gene mal2 at 204 bp as shown in Figure 3.1 plate D.

**Figure 3.1:**
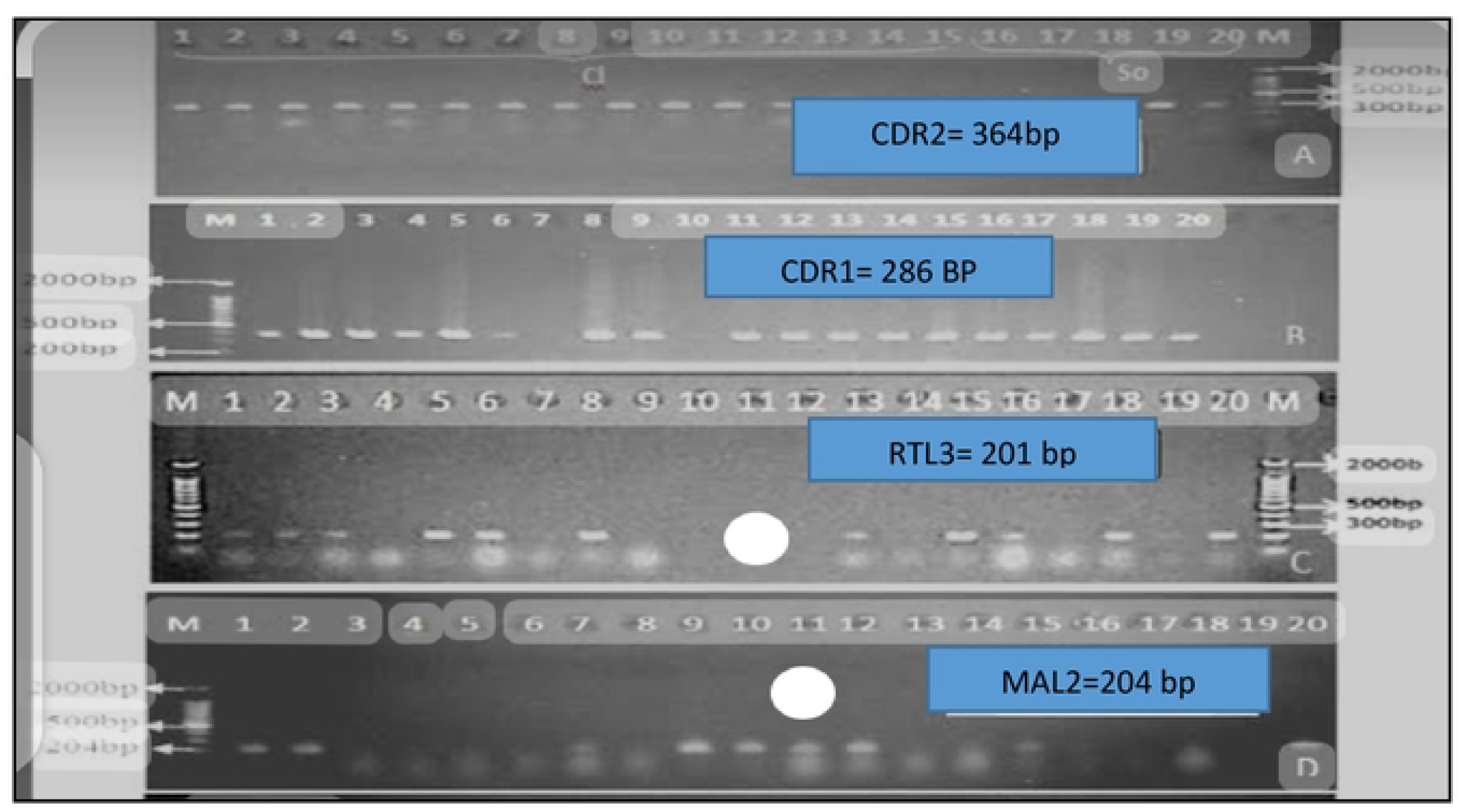
The Gel electrophoresis outcome for confirmation of various resistance genes extracted from the 20 isolates of C.*albicans* isolates **Plate A:** representing the cdr2gene in clinical *C. albicans* isolate. **Plate B:** Representing cdrl genes in clinical *C. albicans* isolate. **Plate C:** Representing rtl3 genes in clinical *C. albicans* isolate and Plate D: Representing mal2 genes in clinical *C. albicans* isolate. Lane 1 (MCH/05/20 isolate); Lane 2 (MCH/06/20 isolate); Lane 3 (MCH/07/20 isolate); Lane 4 (MCH/13/20 isolate); Lane 5 (MCH/14/20 isolate); Lane 6 (MCH/16/20 isolate); Lane 7 (MCH/33/20 isolate); Lane 8 (MCH/34/20 isolate); Lane 9 (MCH/36/20 isolate); Lane 10 (MCH/37/20 isolate); Lane 11 (MCH/38/20 isolate); Lane 12 (MCH/39/20 isolate); Lane 13 (MCH/47/20 isolate); Lane 14 (MCH/47/20 isolate^*^); Lane 15 (MCH/62/20 isolate); Lane 16 (MCH/67/20 isolate); Lane 17 (MCH/74/20 isolate); Lane 18 (MCH/75/20 isolate); Lane 19(MCH/82/20 isolate); Lane 20 (MCH/123/20 isolate); Lane Ml for plate A represents 2 kb DNA Size Marker-Hyper ladder I *cdrl* band 289 bp; lane Ml for plate B represents 2 kb DNA Size Marker-Hyper ladder I *cdr2 band* 364 bp, lane M1 for plate C represents 2 kb DNA Size Marker-Hyper ladder I, *rtl3 band* 201 bp and lane M1 for plate D represents 2 kb DNA Size Marker-Hyper ladder I, *mal2 band* 204 bp.

. The summary on the frequencies of the resistant genes profiling of *C. albicans* isolates is presented in table 3.7 below.

**Table 3.7:** Analysis of pathogenic and antifungal resistance genes by PCR in *C. albicans* isolates

| Target Gene | Positive N (%) | Negative N (%) | Total |
| --- | --- | --- | --- |
| • CDR1 | 8 (40%) | 12 (60%) | 20 |
| • CDR2 | 5 (25%) | 15 (75%) | 20 |
| • RTL3 | 4 (20%) | 16 (80%) | 20 |
| • MAL2 | 3 (15%) | 17 (85%) | 20 |

### vii. Biofilm formation inhibitory effects of selected antifungals against the *C. albicans* strains

Biofilm formation inhibitory effects of selected antifungal drugs against the *C. albicans* strains that were resistant drugs towards the four isolates (Fluconazole, Panosoconazole, Itraconazole, Amphotericin B, Nystatin and Clotrimazole) (36) were used as treatments for the antibiofilm formation assay in a 96-well microtiter plate. The results showed that the biofilm formation inhibitory effects of the various concentrations (0.5, 0.25, 0.125, 0.0625 and 0.03125mgml−1) were significantly lower than that of the positive control, an indication that biofilm formation was inhibited at these concentrations (Figs. 3.2 - 3.5). As much as such inhibitory effects were recorded, these findings clearly demonstrate that the four isolates that proved to be resistant to commonly used antifungal have the ability of forming biofilms.

**Fig. 3.2:**
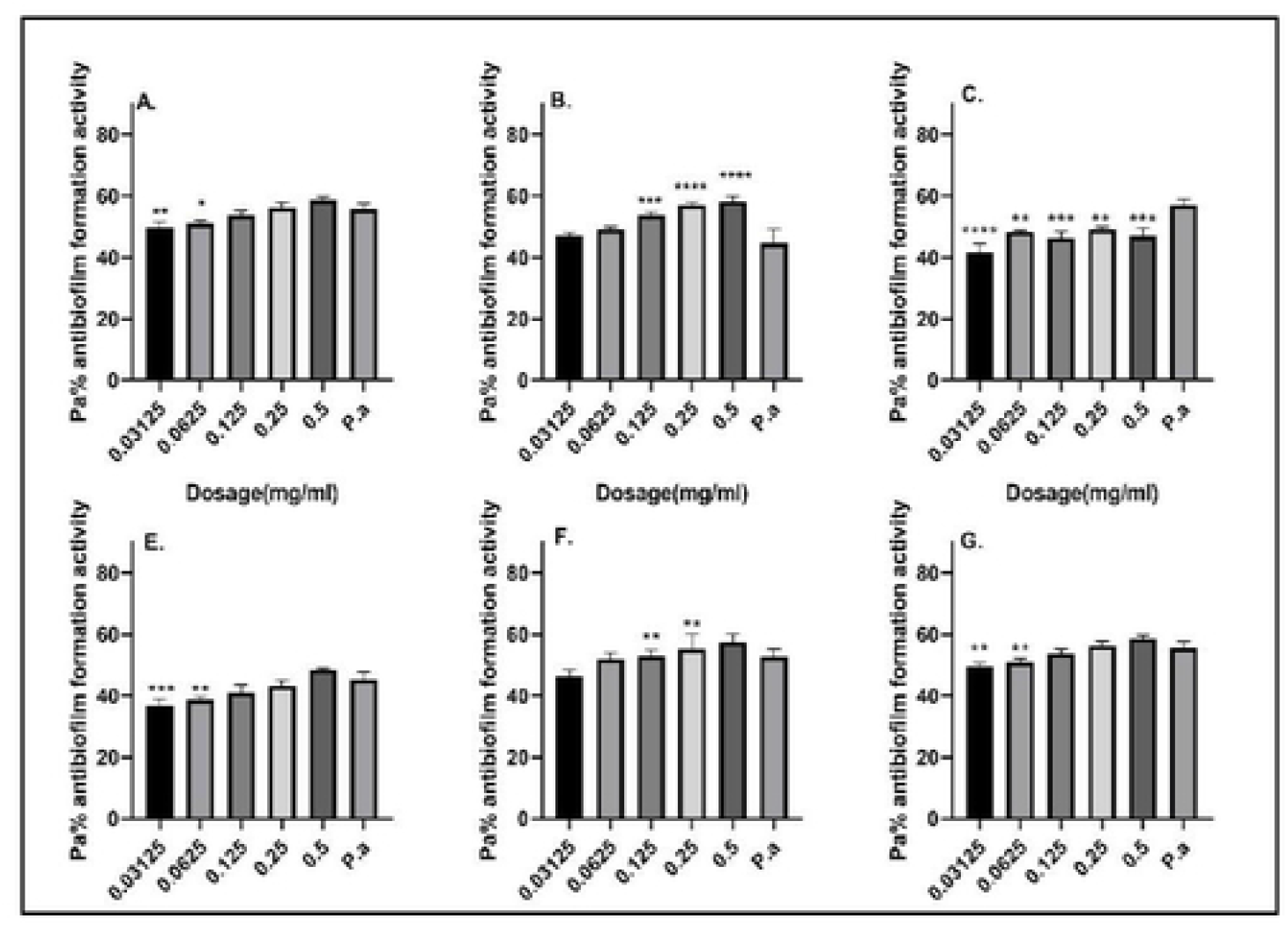
Antibiofilm formation activity against isolate MCH/05/20 of *C. albicans* against various antifungals: (a) Fluconazole, (b) Panosoconazole, (c) Itraconazole, (d) Amphotericin B, (e) Nystatin and (f) Clotrimazole; P. a= *C. albicans* - ATCC 10231–Positive control *(n=*3, ANOVA Dunnett’s multiple comparisons test; \**P*=0.05; \*\**P*=0.01; \*\*\**P*=0.001; \*\*\*\**P*=0

Antibiofilm formation activity against isolate MCH/05/20 of *C. albicans* for various antifungals was low (Fig. 3.2). Interestingly, the concentrations of 0.5, 0.25, 0.125, 0.0625 and 0.03125 mg ml^−1^ were able to inhibit biofilm formation much more as compared with the positive control. A much biofilm inhibitory effects were observed with Nystatin as compared to other antifungals. Additionally, more inhibitory activities were observed at lower dosages for this isolate as compared to higher dosages of the antifungals used as treatments.

Antibiofilm formation activity against isolate MCH/16/20 of *C. albicans* against various antifungals was seen in all the antifungals, with no significance differences on Fluconazole at all the concentration as compared with the positive control (Fig. 3.3). The rest of the antifungals produced significant antibiofilm activity. Amphotericin B, Nystatin and Clotrimazole also showed significant inhibitory activity against this isolate MCH/16/20 at lower concentrations. Also, for this isolate much inhibitory effects were observed at lower dosages of the antifungals used except for Itraconazole.

**Fig. 3.3:**
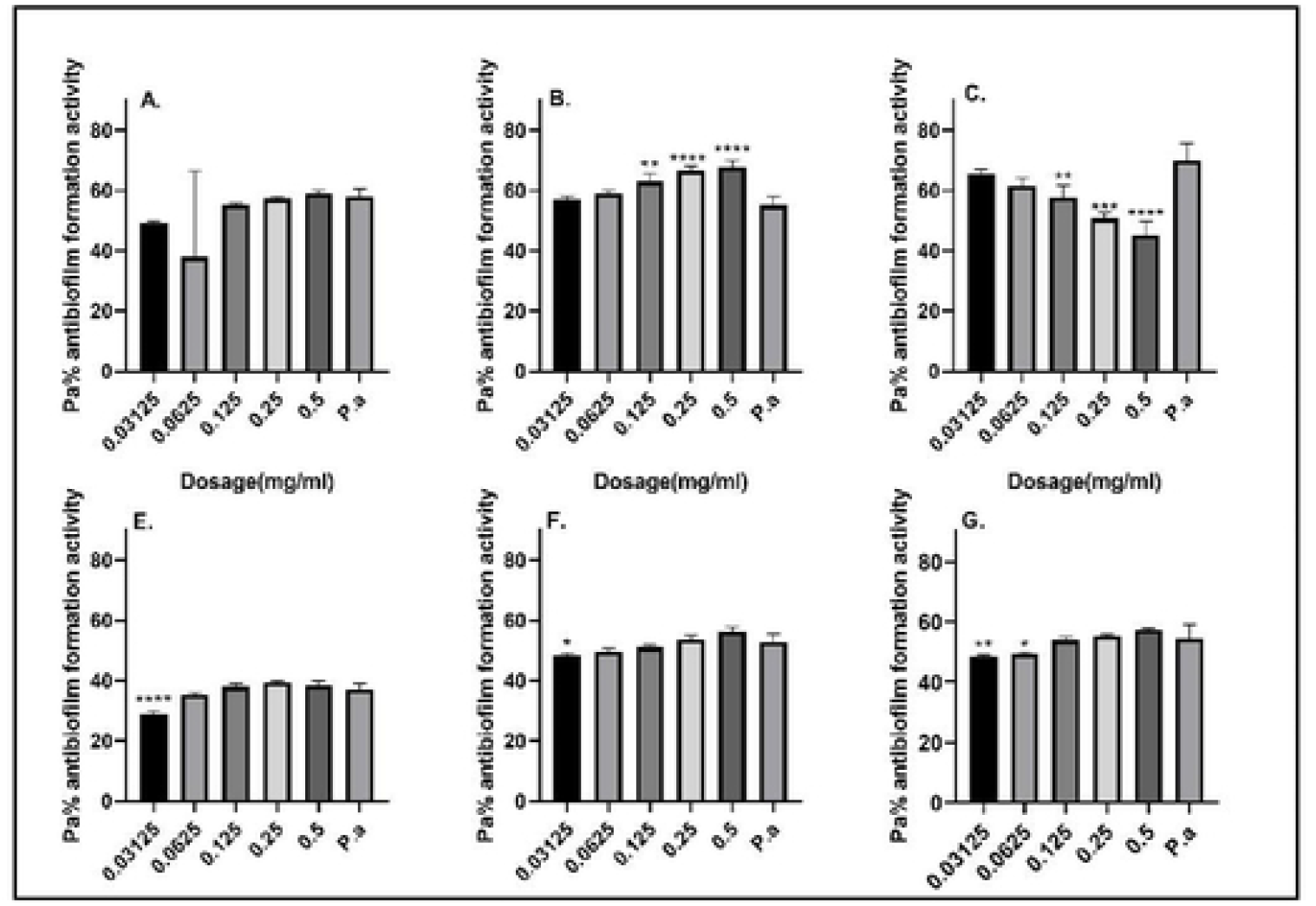
Antibiofilm formation activity against isolate MCH/16/20 of *C. albicans* against various antifungals: (a) Fluconazole, (b) Panosoconazole, (c) Itraconazole, (d) Amphotericin B, € Nystatin and (f) Clotrimazole; P. a= *C. albicans* - ATCC 10231–Positive control *(n=*3, ANOVA Dunnett’s multiple comparisons test; \**P*=0.05; \*\**P*=0.01; \*\*\**P*=0.001; \*\*\*\**P*=0.0001).

Isolate MCH/47/20 was more prone to the inhibitory effects of the various dosages of the various antifungals as compared to other isolates. Significant differences were also observed in most of the antifungals used compared with the positive control (Fig. 3.4). Also, for this isolate much inhibitory effects were observed at lower dosages as compared to higher dosages of the antifungals used except for Itraconazole with Nystatin producing more inhibitory effects at 0.03125 mg/ml.

**Fig. 3.4:**
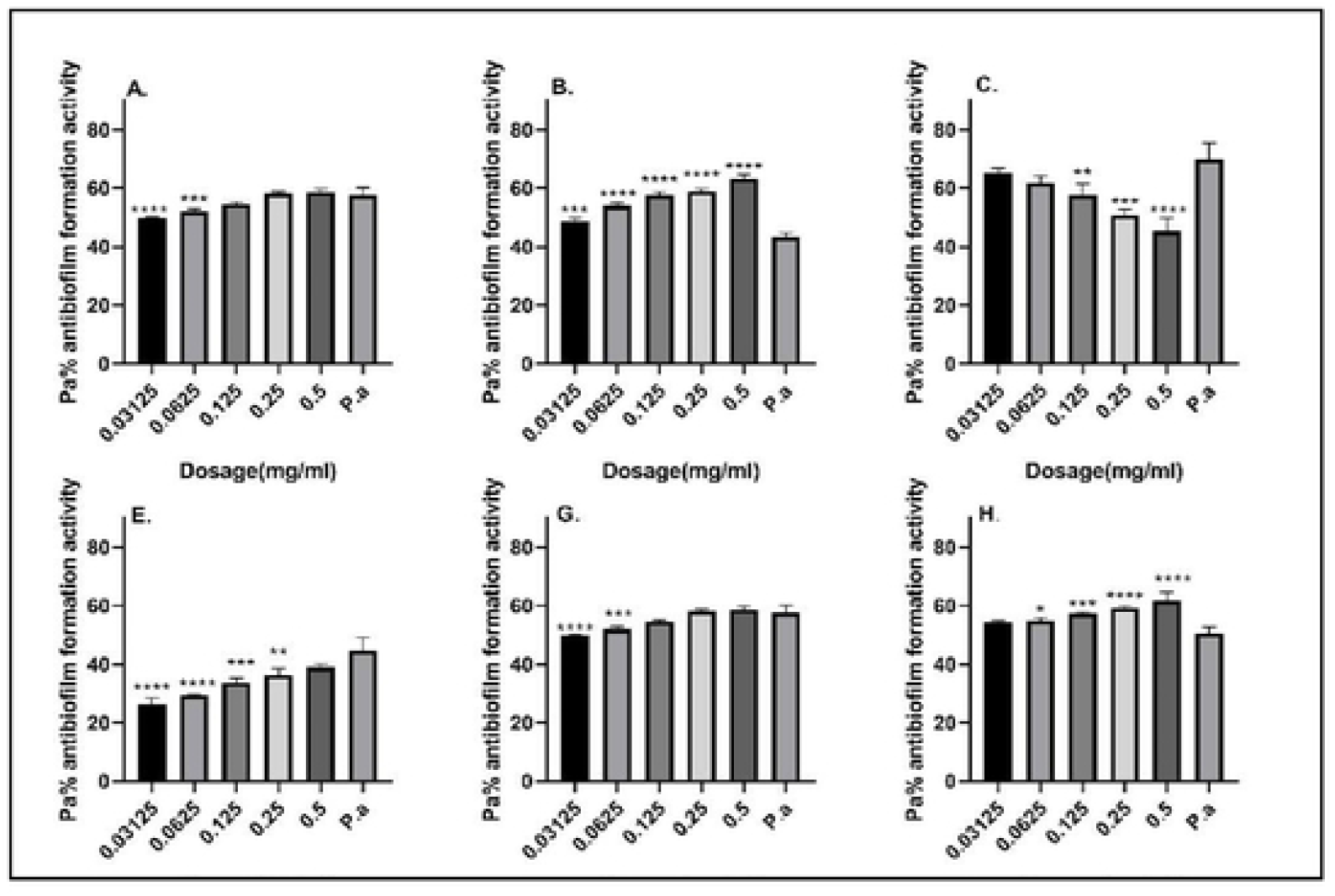
Antibiofilm formation activity against isolate MCH/47/20 of *C. albicans* against various antifungals: (a) Fluconazole, (b) Panosoconazole, (c) Itraconazole, (d) Amphotericin B, (e) Nystatin and (f) Clotrimazole; P. a= *C. albicans* - ATCC 10231–Positive control *(n*=3, ANOVA Dunnett’s multiple comparisons test; \**P*=0.05; \*\**P*=0.01; \*\*\**P*=0.001; \*\*\*\**P*=0.0001).

Antibiofilm formation activity against isolate MCH/75/20 of *C. albicans* against various antifungals was seen in all the antifungals (Fig. 3.5), with some significant differences at various dosages. Panosoconazole not have good inhibitory effects against isolate MCH/75/20. On the other hand, Nystatin showed a reverse activity with high inhibitory effects being observed at 0.03125 mg ml^−1^ as compared with higher concentrations. Similar findings were also observed at lower dosages as compared to higher dosages of the other antifungals used except for Itraconazole.

**Fig. 3.5:**
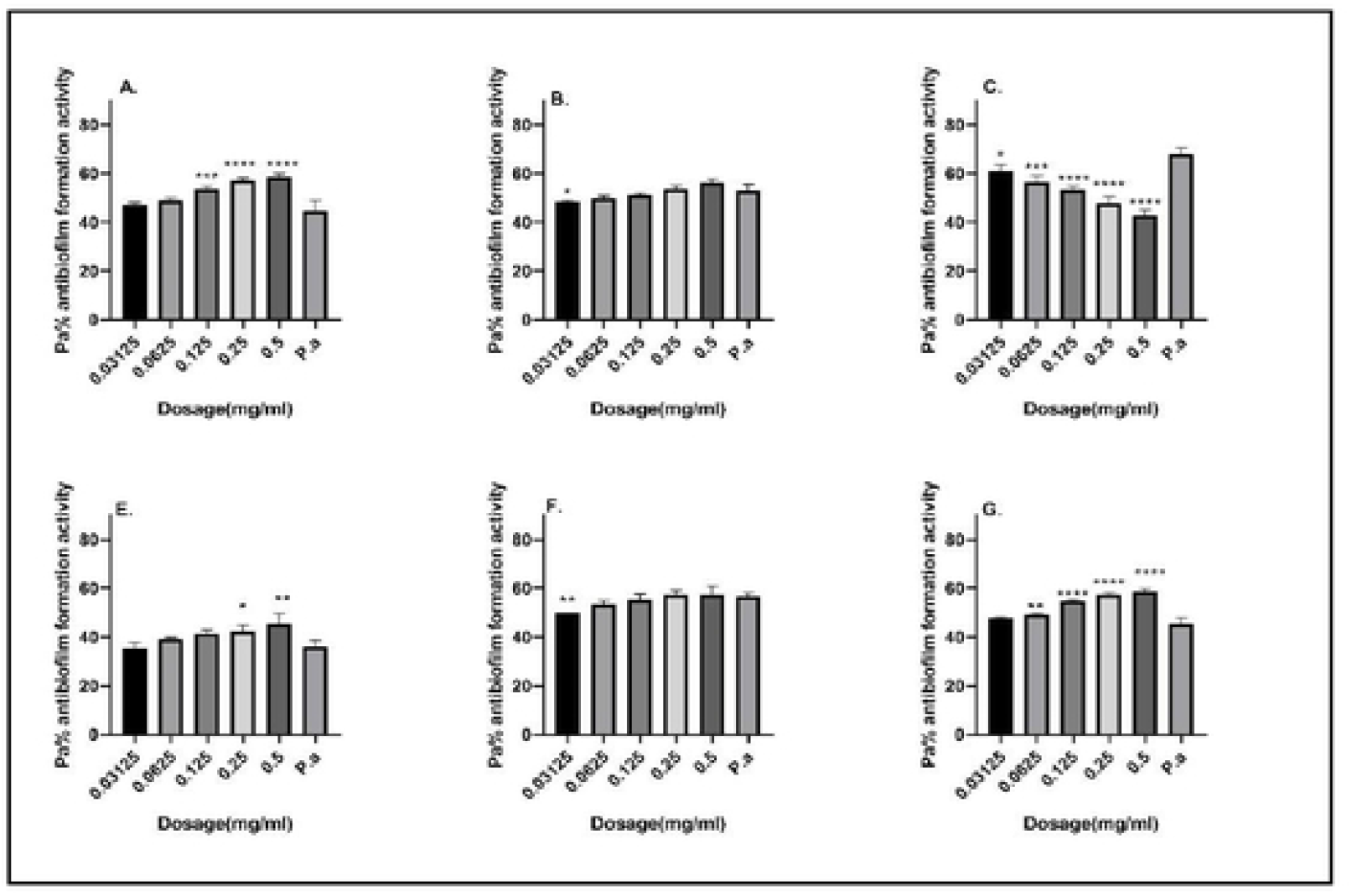
Antibiofilm formation activity against isolate MCH/75/20 of *C. albicans* against various antifungals: (a) Fluconazole, (b) Panosoconazole, (c) Itraconazole, (d) Amphotericin B, (e) Nystatin and (f) Clotrimazole; PC= *C. albicans* - ATCC 10231 – Positive control *(n=* 3, ANOVA Dunnett’s multiple comparisons test; \**P*=0.05;\*\**P*=0.01;\*\*\**P*=0.001; \*\*\*\**P*=0.0001).

## DISCUSSION

Oropharyngeal candidiasis is the most common opportunistic infection among HIV-seropositive patients and in those with AIDS, and it represents a major treatment challenge. Hence, it is recommended to determine the isolate involved in the infection and its antifungal susceptibility. This study showed that 51 samples were from men and 130 were women, resulting in the ratio of 1.3 cases in women for every 1 case in men. These findings lead us to reflect on whether the incidence of AIDS in women is increasing and equating with the incidence in males. Additionally, the disparities in terms of gender could be attributed to the high rate of the incidence (24). Our findings however do not concur with findings from a study done at Brazil, and was reported by Bulletin in 2011, which showed that in 2011, the ratio of 1.7 cases in men for every 1 case in women was greatly diminished when compared to data from 2008, when there were about six cases of AIDS in men for every case in women (24). In the present study, we also found out that the most prevalent age group with candidiasis was 16-25 years, followed by the 26–35 years age group. These findings show similarity to a previous study done in Brazil (24) and other countries (25). The findings on the age group with most prevalent Candida sp. Isolates could be attributed to high rate of sexual practice among the women (31).

During the study period 46 (25.4%) out of the 181 study participants were positive for Candida ssp. culture. This result is similar to that found in other studies, which showed that 20–35% of the patients develop one or more fungal infections during their illness (30, 31,32). In this study, we observed that *C. albicans* was the most prevalent species at 20(43.5%), followed by *C. krusei* 8 (17.4%), *C. tropicalis* 6 (13.0%), *C. glabrata* 4 (8.7%), *C. famata* and *C. parapsilosis* at 3 (6.5%) both, and lastly *C. guilliermondii* 2 (4.3%) which is in accordance with other studies (33,34). These findings could be attributed to the fact that *C. albicans* is sometimes a normal flora in our systems and hence the likelihood of isolating it in higher numbers could be higher. Other studies have also documented the same findings. For instance, in a study done at India they did document that *C. albicans* was the most prevalent species at 30(48.5%) (32).

In this study, we identified a case of a patient with four *Candida spp*. isolates that were identified in a single clinical sample; one of the species was *C. glabrata*, which was resistant to both Fluconazole and Amphotericin, and the other was *C. krusei*, which was resistant to Fluconazole. This finding is relevant because Fluconazole is the drug of choice for candidiasis treatment in AIDS patients although it has a fungistatic action (35,36), and both Fluconazole and Amphotericin were being used by some of the patients who participated in the research. Four patients had double colonization, and one of them had colonization by *C. krusei* resistant to Fluconazole. The coexistence of various species in the same clinical specimen has also been reported in other studies (37,32). The occurrence of oropharyngeal or esophageal candidiasis is recognized as an indicator of immune suppression and is most often observed in patients with CD4 T lymphocyte (CD4) cell counts **<200 cells/mm**^**3**^, with esophageal disease typically occurring at lower CD4 counts than oropharyngeal disease. (37).

In this study, all isolates of *C. albicans* (n D 29), *C. tropicalis* (n D 6), *C. parapsilosis* (n D 2), and *C. guilliermondii* (n D 2) showed sensitivity to all of the antifungal drugs tested, which is consistent with the general pattern of susceptibility of the NCLS M-27 method and with the results of other studies (38).

Ninety percent of the *C. albicans* studied were sensitive to Clotrimazole with regard to 5-FC, a high sensitivity was observed in this study probably because this antifungal action primarily by damaging the permeability barrier in the fungal cytoplasmic membrane (42). Clotrimazole thereby inhibits the biosynthesis of ergosterol in a concentration-dependent manner by inhibiting the demethylation of 14 alpha lanosterol (43)., followed by 85% isolates to Panosoconazole,80% isolates to Fluconazole which interacts with 14-demethylase, a cytochrome P-450 enzyme responsible for catalyzing the conversion of lanosterol to ergosterol (42). As ergosterol forms a critical part of the fungal cell membrane, fluconazole inhibits the synthesis of ergosterol to increase cellular permeability., 70% isolates to both Amphotericin B which binds to ergosterol in the fungal cell membrane, which leads to the formation of pores, ion leakage and ultimately fungal cell death and Nystatin that acts by binding to sterols in the plasma membranes of fungi causing the cells to leak, eventually leading to fungal cell death (43). Lastly 50 % of the isolate show sensitivity to Itraconazol which inhibits the fungal-mediated synthesis of ergosterol, via inhibition of lanosterol 14α-demethylase (44). These results are similar to those observed by other researchers (38,25). Resistance to Itraconazole was observe in 10 (50%) isolants of *C. albicans* followed by 6 (30%) to both Amphotericin and Nystatin, 4 (20%) isolants to Fluconazole, 3 (15%) isolants to Panosoconazole and lastly 2 (10%) isolants to Clotrimazole was observed in this study. The results of this study agree with several studies that demonstrated the innate resistance of *C. albicans* to Fluconazole which is the commonly used antifungal among those living with HIV/AIDs (31, 25, 38, 32, 43).

Further, the study investigated the ability of these test strains to produce various virulence factors, which may play a role in their pathogenicity. Among the virulence traits examined include enzymes like proteinase, phospholipase, capsulase, coagulase and haemolysin on the four isolates, which were found to be resistant to all antifungal revealed that 3/4 (75.0 %) of these isolates of *C. albicans* produced proteinase enzyme, these findings concurs with the findings of a previous study (44), which showed that most isolates were protease positive, and that protease enzyme have limited effect on the pathogenesis of this antifungals. Findings from the current study also agree with a study that documented that all isolates had the ability for protease production (45). Proteases produced by *C. albicans* have a critical role in pathogenicity, as they are responsible for hydrolysis of several physiologically important proteins such as mucin, fibronectin and lactoferrin [46]. It could also proteolytically activate genes and haemolysin, hence making this pathogen more virulent [47].

It was also confirmed that two out four isolates (50.0%) produce phospholipases, these findings further concur with previous study findings were out of 20 isolates, 10 (50 %) isolates were found to have phospholipase production potential (48). Our findings are also in tandem with (49) study findings for phospholipase presence. Phospholipases are lipolytic enzymes that hydrolyze phospholipid substrates at specific ester bonds. Phospholipases are widespread in nature and play very diverse roles from aggression in snake venom to signal transduction, lipid mediator production, and metabolite digestion in humans (50). Further findings also indicate that out of the four isolates two of them (50.0%) had the ability to produce coagulase and only one isolate (25.0%) produced capsulase. Our findings also concur with other studies, which showed that 50% of isolates obtained in the study had the ability to produce coagulase and only 20% produced capsulase which protect fungal from engulfment by host macrophages (50). Coagulase enzymes catalyze the hydrolysis of the ester bonds of triacylglycerols and may have a critical role in *C. albicans* pathogenicity or nutrition acquisition. The production of an excess amount of coagulase allows antifungal to penetrate fatty tissue with the consequent formation of abscesses (51). Therefore, the production of these enzymes by the isolates may reflect the presence of genetic organization of a discrete genetic element, which encodes three genes responsible to produce proteinase, coagulase and phospholipase. This organization could be a possible part of pathogenic island, encoding a product capable of damaging host cells and being involved in nutrient acquisition (51).

Lastly, it was determined that all the four isolates were able to produce the haemolysin by haemolysing the sheep red blood cells causing beta (β) haemolysis which became resistance to host defense, tissue damage, and lethality, either by direct action or by stimulation of inflammatory mediators and signal transduction pathways. Haemolysin therefore was the most produced virulence factor by these isolates (100%) and it was followed by proteinase (75.0%), followed by phospholipase and coagulase at (50%) and lastly capsulase (25.0%). Out of the four isolates one isolates (MCH/16/20) produced all the virulence traits studied, one isolates (MCH/05/20) produced at least three virulence traits and two isolates (MCH/75/20, MCH/47/20) produced at least one of the virulence traits. This finding concurs with another study which reported nearly values as in this our report which reveals that out of the 10 isolates all produced proteinase, phospholipase and coagulase while only 2 (20%) were not able to produce capsulase (52).

From this study, it was revealed that four isolates that had shown resistance to common used antifungals indeed possess various factors that enable them either to be resistant or induce various infections in humans, which leads to alteration of drug efflux which is one of the prominent mechanisms of resistance in fungi. The *C. albicans CDR1* gene is a homolog of *S. cerevisiae* PDR5, which encodes a multidrug efflux pump, and *CDR1* is the gene most often associated with energy-dependent drug efflux in FLU-resistant clinical isolates [52]. The <u>MDR</u> phenotype in *C. albicans* has previously been shown to be linked to proteins encoded by *CDR1, CDR2, RTL3* and *MAL2* genes. These proteins act as membrane-localized efflux pumps that pump drugs from the fungal cells [43]. Some of these traits are controlled at the gene level and as such, they can be passed on from one fungal cell to another through the conjugation or otherwise (27). On analysis of the resistant genes eight (40.0 %) isolates possessed the gene *cdr1*and 5 (25.0 %) harbored the *cdr2* gene. It was also deduced that a variety of pathogenic and antifungal resistance genes like 4 (20.0 %) of the isolates harbored the gene encoded by *rtl3* gene. The minority, 3 (15.0 %) were confirmed to possess the gene mal2. This finding concurs with another study which reported nearly values as in this our report (32).

Some studies have also suggested that Fluconazole and Clotrimazole do inhibit growth at high concentrations (40) but from our study, it was deduced that they inhibit biofilm formation at lower concentrations. The same scenario was also observed in most antifungals like Panosoconazole, Itraconazole, Amphotericin and Nystatin and others with such ‘Goldilocks’ effect. A possible explanation to the less activity observed at greater doses could be associated to the aggregation effects of the antifungal agent at site of entry into fungal cell especially at high dosages something that is not observed at lower dosages. It is likely that aggregation may favors biofilm formation as antifungal agent struggle to reach at the point of action and hence the fungi will continue to thrive and hence form more biofilms [47]. This finding agrees with the previous studies on biofilm inhibitions by [48], which showed higher biofilm inhibitory at lower dosage concentration against the positive control. However, it did not concur with the previous study on biofilm inhibition by [49], which showed that Fluconazole growth inhibitory at high concentration but also biofilm inhibitory at high concentration as compared with positive control [49]. On the other hand, Panosoconazole, Itraconazole, Amphotericin and Nystatin were found to inhibit biofilm formation at higher concentration in some isolates and against the positive control. However, it should be noted that they did not fully inhibit biofilm formation ability of the test isolates a clear indication that proper antifungals should be used in management of conditions caused by these isolates.

## CONCLUSION

*C. albicans* was resistant to Itraconazole, Amphotericin B, and Nystatin antifungal, drugs commonly used in the management on HIV/AIDs patient on care in Muhoroni County Hospital. From this study, it can also be concluded that the clinical isolates at Muhoroni county hospital had resistant genes and produced various virulence factors studied; proteinase, phospholipase, capsulase, coagulase and haemolysin. As much as inhibitory effects were recorded, the study findings clearly demonstrate that the isolates were resistant to commonly used antifungal and they do form biofilms. These findings, therefore, add value to the facility on management of the patient who are on care in the facility. Taken together, there is a need to carry out regular surveillance on antifungal drug resistance during outbreaks.

## Acknowledgements

We thank Muhoroni County Hospital for provision of space for this study. Authors also thank JOOTRH Laboratories-Kisumu, Kenya for providing laboratory space and other resources used this study lastly, we would like to thank all patient who accept to participate by providing us with the sample for analysis.

## Author contributions

All authors contributed equally to this work.

## Conflicts of interest

The authors declare that there are no conflicts of interest.

## Notes

### Competing Interest Statement

The authors have declared no competing interest.

## Reference

1. Prescott KA, Harley JM and Klein DA. Microbiology.7th edition. McGraw-Hill. Publication. New York USA. Resistant candidiasis. AIDS RES. HUM. Retroviruses; 2008; 10: 925–929.

2. White TC, Marr KA, and Bowden RA. Clinical, cellular, and molecular factors that contribute to antifungal drug resistance. Clinical Microbiol Reviews 1998; 11(2): 382–402.

3. Wisplinghoff H, Seifert H, Wenzel RP & Edmond MB. Inflammatory response and clinical course of adult patients with nosocomial bloodstream infections caused by Candida spp. Clin Microbiol Infect 2006; 12: 170–177.

4. Vincent JL, Rello J, Marshall J, Silva E, Anzueto A, Martin CD, Moreno R, Lipman J, Gomersall C & other authors. International study of the prevalence and outcomes of infection in intensive care units. JAMA 2009; 302:2323–2329.

5. Vidigal PG & Svidzinski TIE. Yeasts in the urinary and respiratory tracts: is it a fungal infection or not? J Bras Patol Med Lab 2009; 45: 55–64.

6. Li, Y. Y., Chen, W. Y., Li, X., Li, H. B., Li, H. Q., Wang, L., et al. (2013). Asymptomatic oral yeast carriage and antifungal susceptibility profile of HIV infected patients in Kunming, Yunnan Province of China. BMC Infect. Dis. 13:46–53. doi: 10.1186/1471-2334-13-46 Centers for Disease Control and Prevention. Candidiasis. [<http://www.cdc.gov/fungal/diseases/candidiasis/index.html/>. DOI: 10.12865/CHSJ.42.02.08

7. Pfaller MA, Diekema DJ. 2007. Epidemiology of Invasive Candidiasis: A Persistent Public Health Problem. Virulence. (2): 119–128.

8. Public Health Agency of Canada. Candida albicans - Material Safety Data Sheets. [<http://www.phac-aspc.gc.ca/lab-bio/res/psdsftss/msds30e-eng.php>.

9. NCCLS. Method for Antifungal Disk Diffusion Susceptibility Testing of Yeasts; Approved Guideline. NCCLS document M44-A [ISBN 156238-532-1]. NCCLS, 940 West Valley Road, Suite 1400, Wayne, Pennsylvania 19087-1898 USA, 2004. Lai CC, Wang CY, Liu WL, Huang YT & Hsueh

10. Awuor SO, Omwenga EO, Daud II. Geographical distribution and antibiotics susceptibility patterns of toxigenic Vibrio cholerae isolates from Kisumu County, Kenya. African Journal of Primary Health Care & Family Medicine. 2020;12(1).

11. Arendrup MC, Cuenca-Estrella M, Donnelly JP, et al. EUCAST technical notes on Posaconazole. Clin Microbiol Infect 2011; 17 (11): E16–E17.

12. Lass-Flörl C, Arendrup MC, Rodriguez-Tudela J-L, et al. EU-CAST technical notes on Amphotericin B. Clin Microbiol Infect 2011; 17 (12): E27–E29.

13. Owotade FJ, Patel M, Ralephenya TRMD, Vergotine G. Oral Candida colonization in HIV positive women: associated factors and changes with antiretroviral therapy. J Med Microbiol 2013; 62 (Pt 1): 126–32.

14. Hospenthal, D. R., Murray, C. K., and Rinaldi, M. G. (2004). The role of antifungal susceptibility testing in the therapy of candidiasis. Diagn. Microbiol. Infect. Dis. 48, 153– 160. doi: 10.1016/j.diagmicrobio.2003.10.003

15. Borghi, E., Iatta, R., Sciota, R., Biassoni, C., Cuna, T., Montagna, M. T., et al. (2010). Comparative evaluation of the Vitek 2 yeast susceptibility test and CLSI broth microdilution reference method for testing antifungal susceptibility of invasive fungal isolates in Italy: the GISIA 3 study. J. Clin. Microbiol. 48, 3153–3157. doi: 10.1128/JCM.00952-10

16. Redding SW, Zellars RC, Kirkpatrick WR, et al. Epidemiology of oropharyngeal Candida colonization and infection in patients receiving radiation for head and neck cancer. J Clin Microbiol 1999; 37(12): 3896–900.

17. Perlin, DS. Antifungal drug resistance in developing countries. In: Sosa AJ, Byarugaba DK, Amabile-Cuevas CF, Hsueh PR, Kariuki S, Okeke IN, eds. Springer New York Dordrecht Heidelberg London; 2010, p. 137–56.

18. Kaur, R., Dhakad, M. S., Goyal, R., Haque, A., and Mukhopadhyay, G. (2016). Identification and Antifungal susceptibility testing of Candida species: a comparison of Vitek-2 system with conventional and molecular methods. J. Glob. Infect. Dis. 8, 139–146. doi: 10.4103/0974-777X.192969

19. Patil, S., Rao, R. S., Majumdar, B., and Anil, S. (2015). Clinical appearance of oral Candida infection and therapeutic strategies. Front. Microbiol. 6:1391. doi: 10.3389/fmicb.2015.01

20. Peano, A., Pasquetti, M., Tizzani, P., Chiavassa, E., Guillot, J., & Johnson, E. (2017). Methodological issues in antifungal susceptibility testing of Malassezia pachydermatis. Journal of Fungi, 3(3), 37.

21. Hamza OJM, Matee MIN, Moshi MJ, et al. Species distribution and in vitro antifungal susceptibility of oral yeast isolates from Tanzanian HIV-infected patients with primary and recurrent oropharyngeal candidiasis. BMC Microbiol 2008; 8:135. Doi:10.1186/1471-2180-8-135.

22. Rodríguez-Cerdeira, C., Gregorio, M. C., Molares-Vila, A., López-Barcenas, A., Fabbrocini, G., Bardhi, B., & Hernandez-Castro, R. (2018). Biofilms and vulvovaginal candidiasis. Colloids and Surfaces B: Biointerfaces.

23. Zaranza, A.V.; Morais, F.C.; do Carmo, M.S.; de Mendonça Marques, A.; Andrade-Monteiro, C.; Ferro, T.F.; Monteiro-Neto, V.; Figueiredo, P.d.M.S (2013). Antimicrobial susceptibility, biofilm production and adhesion to HEp-2 cells of Pseudomonas aeruginosa strains isolated from clinical samples. J. Biomater. Nanobiotechnol. 4, 98–106.

24. Hinrichsen, S. L., Falcao, E., Vilella, T. A., Colombo, A. L., Nucci, M., Moura, L., et al. (2008). Candidemia in a tertiary hospital in northeastern Brazil. Rev. Soc. Bras. Med. Trop. 41, 394–398. doi: 10.1590/S0037-86822008000400014

25. Hamza, O. J., Matee, M. I., Moshi, M. J., Simon, E. N., Mugusi, F., Mikx, F. H., et al. (2008). Species distribution and in vitro antifungal susceptibility of oral yeast isolates from Tanzanian HIV-infected patients with primary and recurrent oropharyngeal candidiasis. BMC Microbiol. 8:135. doi: 10.1186/1471-2180-8-135

26. Clinical and Laboratory Standards Institute [CLSI] (2022). M27-A3 Reference Method for Broth Dilution Antifungal Susceptibility Testing of Yeast; Approved Standard, 32rd Edn, Wayne, PA: CLSI, 6–12.

27. Bourgeois, N., Dehandschoewercker, L., Bertout, S., Bousquet, P. J., Rispail, P., and Lachaud, L. (2010). Antifungal susceptibility of 205 Candida spp. Isolated primarily during invasive Candidiasis and comparison of the Vitek 2 system with the CLSI broth microdilution and Etest methods. J. Clin. Microbiol. 48, 154–161. doi: 10.1128/JCM.01096-09

28. Cuenca-Estrella, M., Gomez-Lopez, A., Alastruey-Izquierdo, A., Martinez, L. B., Cuesta, I., Buitrago, M. J., et al. (2010). Comparison of the Vitek 2 antifungal system with the clinical and laboratory standards institute (CLSI) and European committee on antimicrobial susceptibility testing (EUCAST) broth microdilution reference methods and with the sensititre yeast one and etest techniques for in vitro detection of antifungal resistance in yeast isolates. J. Clin. Microbiol. 48, 1782–1786. doi: 10.1128/JCM.02316-09

29. Rex, J. H., Walsh, T. J., Sobel, J. D., Filler, S. G., Pappas, P. G., Dismukes, W. E., et al. (2000). Practice guidelines for the treatment of candidiasis. Clin. Infect. Dis. 30, 662–678. doi: 10.1086/313749

30. Campisi, G., Pizzo, G., Milici, M. E., Mancuso, S., and Margiotta, V. (2002). Candidal carriage in the oral cavity of human immunodeficiency virus-infected subjects. Oral Surg. Oral Med. Oral Pathol. Oral Radiol. Endod. 93, 281–286. doi: 10.1067/moe.2002.120804

31. Junqueira, J. C., Vilela, S. F., Rossoni, R. D., Barbosa, J. O., Costa, A. C. B. P., Rasteiro, V. M. C., et al. (2012). Oral colonization by yeasts in HIV-positive patients in Brazil. Rev. Inst. Med. Trop. Sao Paulo 54, 17–24. doi: 10.1590/S0036-46652012000100004

32. Viudes, A., Peman, J., Canton, E., Ubeda, P., Lopez-Ribot, J. L., and Gobernado, M. (2002). Candidemia at a tertiary-care hospital: epidemiology, treatment, clinical outcome and risk factors for death. Eur. J. Clin. Microbiol. Infect. Dis. 21, 767–774. doi: 10.1007/s10096-002-0822-1

33. Colombo, A. L., Nucci, M., Park, B. J., Nouér, S. A., Arthington-Skaggs, B., da Matta, D. A., et al. (2006). Epidemiology of candidemia in Brazil: a nationwide sentinel surveillance of candidemia in eleven medical centers. J. Clin.Microbiol. 44, 2816–2823. doi: 10.1128/JCM.00773-06

34. Siikala, E., Rautemaa, R., Richardson, M., Saxen, H., Bowyer, P., and Sanglard, D. (2010). Persistent Candida albicans colonization and molecular mechanisms of azole resistance in autoimmunepolyendocrinopathy-candidiasis-ectodermal dystrophy (APECED) patients. J. Antimicrob. Chemother. 65, 2505–2513. doi: 10.1093/jac/dkq354

35. Rautemaa, R., and Ramage, G. (2011). Oral candidosis – clinical challenges of a biofilm disease. Crit. Rev. Microbiol. 37, 328–336. doi: 10.3109/1040841X.2011.585606

36. Swinne, D., Watelle, M., Van der Flaes, M., and Nolard, N. (2004). In vitro activities of voriconazole (UK-109, 496), fluconazole, itraconazole and amphotericin B against 132 non-albicans bloodstream yeast isolates (CANARI study). Mycoses 47, 177–183. doi: 10.1111/j.1439-0507.2004.00971.x

37. Pfaller, M. A., Diekema, D. J., Gibbs, D. L., Newell, V. A., Nagy, E., Dobiasova, S., et al. (2008). Candida krusei, a multidrug-resistant opportunistic fungal pathogen: geographic and temporal trends from the ARTEMIS DISK antifungal surveillance program, 2001 to 2005. J. Clin. Microbiol. 46, 515–521. doi: 10.1128/JCM.01915-07

38. Bourgeois, N., Dehandschoewercker, L., Bertout, S., Bousquet, P. J., Rispail, P., and Lachaud, L. (2010). Antifungal susceptibility of 205 Candida spp. Isolated primarily during invasive Candidiasis and comparison of the Vitek 2 system with the CLSI broth microdilution and Etest methods. J. Clin. Microbiol. 48, 154–161. doi: 10.1128/JCM.01096-09

39. Cuenca-Estrella, M., Gomez-Lopez, A., Alastruey-Izquierdo, A., Martinez, L. B., Cuesta, I., Buitrago, M. J., et al. (2010). Comparison of the Vitek 2 antifungal system with the clinical and laboratory standards institute (CLSI) and European committee on antimicrobial susceptibility testing (EUCAST) broth microdilution reference methods and with the sensititre yeast one and etes techniques for in vitro detection of antifungal resistance in yeast isolates. J. Clin. Microbiol. 48, 1782–1786. doi: 10.1128/JCM.02316-09

40. Kaur, R., Dhakad, M. S., Goyal, R., Haque, A., and Mukhopadhyay, G. (2016). Identification and Antifungal susceptibility testing of Candida species: a comparison of Vitek-2 system with conventional and molecular methods. J. Glob. Infect. Dis. 8, 139–146. doi: 10.4103/0974-777X.192969

41. Bremenkamp, R. M., Caris, A. R., Jorge, A. O., Back-Brito, G. N., Mota, A. J., Balducci, I., et al. (2011). Prevalence and antifungal resistance profile of Candida spp. oral isolates from patients with type 1 and 2 diabetes mellitus. Arch. Oral Biol. 56, 549–555. doi: 10.1016/j.archoralbio.2010.11.018

42. Eddouzi J, Lohberger A, Vogne C, Manai M, Sanglard D. Identification and antifungal susceptibility of a large collection of yeast strains isolated in Tunisian hospitals. Med Mycol 2013; 51(7): 737–46.

43. Abrantes PMDS, McArthur CP, Africa CWJ. Multi-drug resistant (MDR) oral Candida species isolated from HIV-positive patients in South Africa and Cameroon. Diagn Microbiol Infect Dis 2014; 79 (2): 222–7.

44. Nweze EI, Ogbonnaya UL. Oral Candida isolates among HIV-infected subjects in Nigeria. J Microbiol Immunol Infect 2011; 44(3): 172–7.

45. Mulu A, Kassu A, Anagaw B, et al. Frequent detection of ‘azole’ resistant Candida species among late presenting AIDS patients in northwest Ethiopia. BMC Infect Dis 2013; 13:82. Doi: 10.1186/1471-2334-13-82.

46. Garcia-Agudo L, Garcia-Martos P, Martos-Canadas J, Aznar-Marin P, Marin-Casanova P, Rodriguez-Iglesias M. Evaluation of the Sensititre Yeast One microdilution method for susceptibility testing of Candida species to anidulafungin, caspofungin, and micafungin. Rev Esp Quimioter 2012; 25(4): 256–60.

47. Badiee P, Alborzi A, Davarpanah MA, Shakiba E. Distributions and antifungal susceptibility of Candida species from mucosal sites in HIV positive patients. Arch Iran Med 2010; 13(4): 282–7.

48. Ma, Boyken L, Hollis RJ, et al. Wild-type MIC distributions and epidemiological cutoff values for posaconazole and voriconazole and Candida spp. as determined by 24-hour CLSI broth microdilution. J Clin Microbiol 2011; 49 (2): 630–7.

49. Hunter KD, Gibson J, Lockhart P, Pithie A, Bagg J. Fluconazole resistant Candida species in the oral flora of fluconazole exposed HIV-positive patients. Oral Surg Oral Med Oral Pathol Oral Radiol Endod 1998; 85 (5): 558–64.

50. Goldman M, Cloud GA, Smedema M, et al. Does long-term itraconazole prophylaxis result in in vitro azole resistance in mucosal Candida albicans isolated from persons with advanced human immunodeficiency virus infection? The National Institute of Allergy and Infectious Diseases Mycoses Study Group. Antimicrobe Agents chemother 2000; 44 (6): 1585–7.

51. Pelletier R, Peter J, Antin C, Gonzalez C, Wood L, Walsh TJ. Emergence of resistance of Candida albicans to clotrimazole in human immunodeficiency virus-infected children: in vitro and clinical correlations. J Clin Microbiol 2000; 38 (4): 1563–8.

52. Rautemaa R, Richardson M, Pfaller M, Perheentupa J, Saxen H. Reduction of fluconazole susceptibility of Candida albicans in APECED patients due to long-term use of ketoconazole and miconazole. Scand J Infect Dis 2008; 40 (11-12): 904–7.

